# Hidden antibiotic resistance fitness costs revealed by GWAS-based epistasis analysis

**DOI:** 10.1101/148825

**Authors:** Maho Yokoyama, Maisem Laabei, Emily Stevens, Leann Bacon, Kate Heesom, Sion Bayliss, Nicola Ooi, Alex J. O’Neill, Ewan Murray, Paul Williams, Anneke Lubben, Shaun Reeksting, Guillaume Meric, Ben Pascoe, Samuel K. Sheppard, Mario Recker, Laurence D. Hurst, Ruth C. Massey

## Abstract

Understanding how multi-drug resistant pathogens evolve is key to identifying means of curtailing their further emergence and dissemination. Fitness costs imposed on bacteria by resistance mechanisms are believed to hamper their dissemination in an antibiotic free environment, however, some have been reported to have little or no cost, which suggests there are few barriers preventing their global spread. One such apparently cost-free resistance mechanism acquired by the major human pathogen *Staphylococcus aureus* is to the clinically important antibiotic mupirocin, which is mediated by mutation of the highly-conserved and essential isoleucyl-tRNA synthethase (*ileS*) gene. In Genome Wide Association Studies (GWAS) on two genetically and geographically distinct MRSA lineages we have found this mutation to be associated with changes in bacterial virulence, driven through epistatic interactions with other loci. Given the potential dual effect of this mutation on both antibiotic resistance and virulence we adopted a proteomic approach and observed pleiotropic effects. This analysis revealed that the activity of the secretory apparatus of the PSM family of cytolytic toxins, the Pmt system, is affected in the mupirocin resistant mutant, which explains why it is less toxic. As an energetically costly activity, this reduction in toxicity masks the fitness costs associated with this resistance mutation, a cost that becomes apparent when toxin production is required. Given the widespread use of this antibiotic, and that this resistance often results from a single nucleotide substitution in the *ileS* gene, these hidden fitness costs provide an explanation for why this resistance mechanism is not more prevalent. This work also demonstrates how population-based genomic analysis of virulence and antibiotic resistance can contribute to uncovering hidden features of the biology of microbial pathogens.

## INTRODUCTION

Antibiotic resistance can evolve in many ways, and frequently incur a fitness cost to the organism^1^ which has to either mutate the target site of the antibiotic, acquire and express a gene encoding an alternative non-susceptible version of the target protein, or acquire and produce an efflux pump that removes the antibiotic before it can attack its target^2^. As antibiotics are most commonly used for short and defined periods of time, resistant bacteria are under selection to reduce these costs to avoid displacement once treatment has finished. In many cases this is achieved through compensatory mutations that allow many resistance mechanisms to be maintained stably populations for long periods of time^3^. However, some antibiotic resistance mechanisms have been reported to incur no detectable fitness costs^4,5,6^, which suggests there are no barriers to their widespread dissemination.

*Staphylococcus aureus* is an example of a major human pathogen^7^ that has become more challenging to treat due to the emergence of antibiotic resistance, with Methicillin-Resistant *S. aureus* (MRSA) being the most notable example^8^. *S. aureus* resides asymptomatically as part of the normal nasal flora of up to 50% of humans^9^, however, this is a significant risk factor for infection^10^, to the extent that carriers are often decolonised using antibiotics such as mupirocin prior to invasive procedures such as surgery or dialysis^11^. Mupirocin is a polyketide antibiotic that is applied as an ointment to eradicate nasal carriage of MRSA in patients at risk of infection^12^. Such decolonisation has been reported to reduce *S. aureus* infections of post-surgical wounds by 58%, of haemodialysis patients by 80% and of peritoneal dialysis patients by 63%^13^.

The molecular target for mupirocin is the bacterial isoleucyl-tRNA synthetase (IleRS), which charges tRNAs with the amino acid isoleucine (lle)^12^. By binding to this enzyme the antibiotic halts protein synthesis, so inhibiting bacterial growth^14^. As a consequence of the widespread use of mupirocin, resistance has emerged where the bacteria have mutated the gene encoding IleRS, *ileS*, resulting in an amino acid substitution (e.g. V588F, encoded by a G to T single nucleotide polymorphism (SNP) at position 1,762 in the *ileS* gene (G1762T)) which alters the structure of the protein’s active site, but retains functionality and renders mupirocin less effective^15^. This confers a low to intermediate level of resistance to the antibiotic^16^.

Alternatively, the bacteria acquire an alternative IleRS, encoded by a *mupA* or *mupB* gene, on a plasmid, which confers a higher level of resistance^17,18^. The prevalence of mupirocin resistance varies widely, with the highest rates associated with patient groups repeatedly exposed to the antibiotic^19,20^. It has also been shown that in countries such as New Zealand and Australia which have previously reported a high prevalence of mupirocin resistance, when restrictions were put in place limiting the use of this antibiotic, the prevalence of resistant strains significantly declined^20^. This suggests that antibiotic exposure is required to maintain selection of this resistance mechanism within a population, despite it being reported as incurring no fitness cost.

The *ileS* gene is highly conserved across the thousands of sequenced *S. aureus* isolates, and many failed attempts to inactivate it suggest its activity is essential to the bacteria. It is therefore surprising that the mutation that confers mupirocin resistance, by altering the structure of the encoded protein, does not appear to affect fitness. Especially considering that the replacement of the valine 588 (V588) with phenylalanine, a much bulkier residue, is likely to fill and distort the Rossman fold of the enzyme, which is responsible for ATP binding activity of this enzyme^4,10^. However, in a recent genome wide association study (GWAS) on the major hospital acquired MRSA clone, ST239, this mutation was significantly associated with differences in the virulence of *S. aureus* isolates^21^. Its effect on toxin secretion, a major aspect of *S. aureus* virulence, was believed to result from epistatic interactions between *ileS* and other polymorphic loci. Upon analysis of a second genetically and geographically distinct collection of clinical isolates of the USA300 lineage we detected this epistasis signal again, and here we characterise the effect this mutation has on both toxin production and bacterial fitness.

## RESULTS and DISCUSSION

### Epistasis between the mupirocin resistance conferring mutation in the *ileS* gene and other loci is associated with the toxicity of the USA300 MRSA lineage

Having previously identified toxicity-associated, epistatic interactions occurring between the mupirocin resistance (mup^R^) conferring mutation in the *ileS* gene (G1762T) and other loci within a collection of ST239 MRSA isolates^21^, we sought to determine whether this was lineage specific. We analysed toxicity and sequence data for a collection of 130 USA300 MRSA isolates^22^, where the SNP conferring mup^R^ resistance emerged for the USA300 isolates as the most dominant toxicity-affecting epistatically-interacting locus (fig. 1), demonstrating the widespread nature of this effect across diverse clonal lineages. See Supp. Table 1 for the list of loci associated with the mup^R^ conferring SNP for both the ST239 and USA300 collections.

**Fig. 1:**
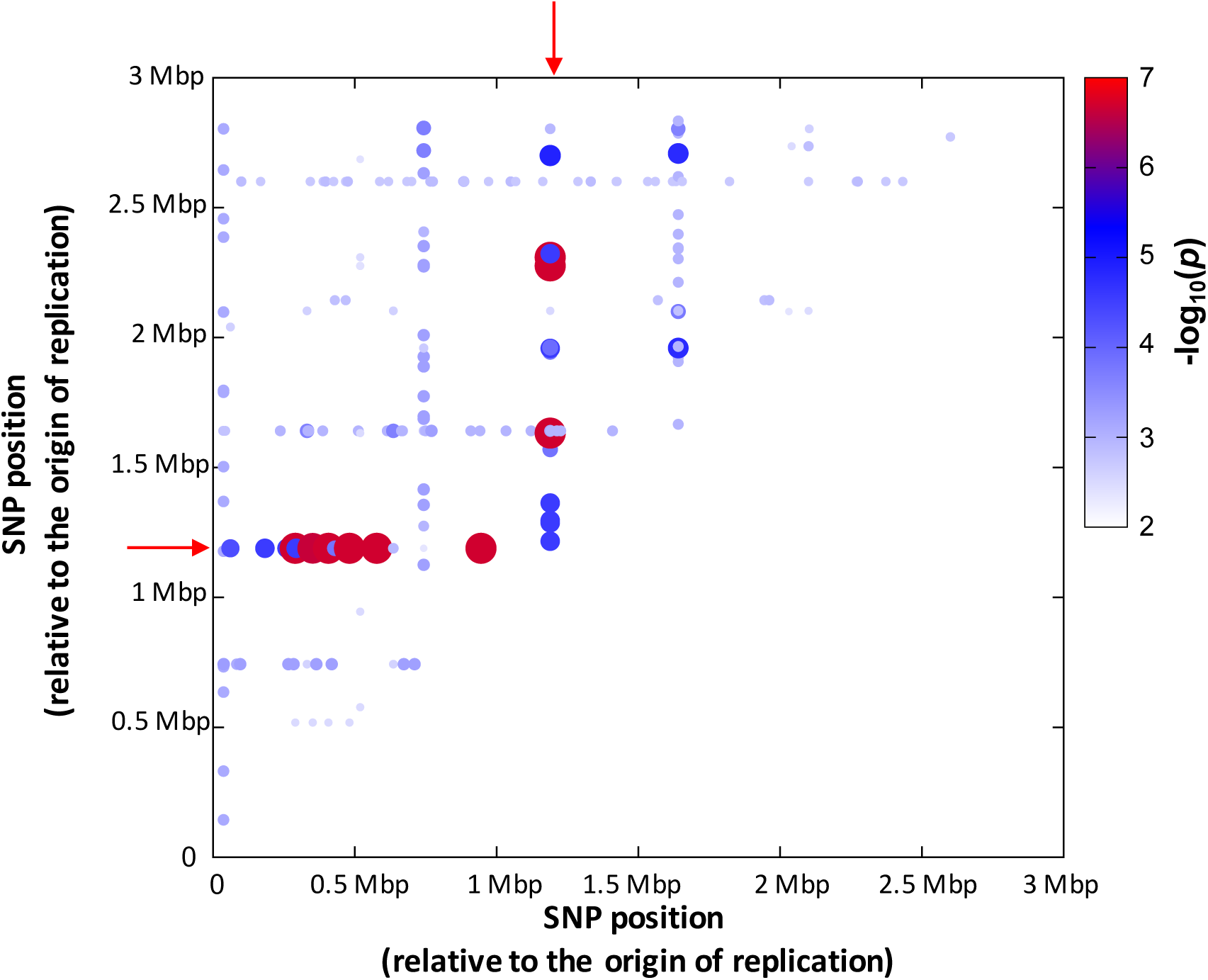
Epistasis between the mupirocin resistance encoding mutation in the *ileS* gene and many other loci is associated with the toxicity of the USA300 lineage of MRSA. This heat map illustrates where specific combinations of the polymorphic site in the *ileS* gene and polymorphic sites elsewhere on the chromosome are associated with the toxicity of individual isolates. The mup^R^ conferring site is indicated on the X and Y axis by the red arrow.

**Table 1:** Toxicity affecting epistatic interactions. An example of a SNP associated by our epistasis analysis to be interacting with the mupirocin resistance conferring SNP to affect toxicity. The mean toxicity of each of the four combinations of the four alleles is presented +/-the 95% confidence intervals.

| Percentage Cell death<br>(+/- 95% confidence) | SAUSA300_0426 (T) | SAUSA300_0426 (C) |
| --- | --- | --- |
| Mupirocin resistant | 45.6% (+/- 5.7) | 4.6% (+/- 2.8) |
| Mupirocin sensitive | 20.65% (+/- 4) | 53% (+/- 9.2) |

To illustrate the epistatic effect on toxicity we selected at random one of the interacting loci and present the mean toxicity of the four combinations of each allele of the two genes (Table 1). The SNP we selected was at position 480640 (relative to the origin of replication) and confers a non-synonymous change in the protein encoded by the open-reading frame with the locus tag SAUSA300_0426, which is described as a conserved hypothetical protein. When in combination with the mup^R^ encoding SNP in *ileS*, strains with the allele of SAUSA300_0426 containing a T at position 480640 are significantly more toxic than those with a C at this site. Whereas in the mupirocin sensitive strains, those with the C at position 480640 are more toxic than those with a T at this site (Table 1). This example illustrates the toxicity affecting epistatic signal detected by our analysis.

### Mupirocin resistance exerts a pleiotropic effect on the *S. aureus* proteome

Across both the ST239 and USA300 collections of isolates, 59 loci associated with the toxicity of *S. aureus* through epistatic interactions with the mup^R^ mutation (Supp. Table 1). To understand how such potential interactions could affect toxicity we examined this list of loci, however, no known toxicity affecting genes were identified, and no loci were common between the two clones. As IleRS is involved in the translation of proteins, and our GWAS data suggests this antibiotic resistance conferring mutation also affects the ability of *S. aureus* to secrete toxins, we hypothesised that this mutation may have pleiotropic effects on protein production, which could explain the observed epistasis. To examine this, we isolated a mup^R^ version of the *S. aureus* laboratory strain SH1000 by plating overnight cultures on agar containing 4*μ*g/ml mupirocin. Mupirocin resistant colonies of the SH1000 strain were recovered, and to confirm that the V588F conferring SNP was present the colonies were sequenced and compared to SH1000. We selected a colony which we have designated MY40, where the only non-synonymous SNP found in this strain was that conferring the V588F change in IleRS, although a small number of non-synonomous SNP differences were detected (Supp. Table 2).

We performed Tandem-Mass-Tagging (TMT) protein mass spectroscopy^23^ on whole cell lysates of the wild type SH1000 and mup^R^ strain MY40. The proteins were extracted from triplicate 18hr cultures in Tryptone Soy Broth (TSB), and of the 3026 open reading frames predicted for the NCTC8325 chromosome (which is the closest reference genome to SH1000), this proteomic approach was able to detect and quantify 1284 proteins, where we used a cut-off of a minimum of a two-fold difference in protein abundance to identify differentially produced proteins. When we compared the proteome of SH1000 and MY40, there were 140 proteins that were differentially produced (Supp. Table 3), which verified our hypothesis that this resistance mutation has pleiotropic effects. However, when we compared this list of differentially produced proteins with our list of loci associated with toxicity through epistasis with the mup^R^ mutation (Supp. Table 1) there was no overlap, suggesting any toxicity affecting interactions between these loci must be indirect.

### Exposure to mupirocin and mupirocin resistance have common effects on protein production

To examine whether the mup^R^ mutation has a similar effect on IleRS activity as the presence of mupirocin has, alongside our proteomic analysis of the the mup^R^ mutant we also analysed lysate of SH1000 exposed to the highest concentrations of mupirocin for which we found no inhibition of growth (10ng/ml). This concentration was selected to avoid any confounding effects differences in growth rates might have on the proteome. Exposure of the wild type SH1000 strain to mupirocin resulted in differential production of 67 proteins when compared to the untreated SH1000 (Supp. Table 4), 18 of which were also different in the mup^R^ mutant, suggesting many common down-steam effects of interfering with the activity of IleRS. These include an increase in the production of several of the proteins involved in Ile biosynthesis (i.e. LeuA, LeuC and IlvA), which suggests that the bacteria are compensating for the interference in IleRS activity due to the mutation and exposure to the antibiotic. Another common expression difference was that several of the iron-regulated surface determinant (Isd) proteins were expressed at lower levels in both the mutant and the mupirocin-exposed wild type when compared with the untreated wild type. As yet we have no explanation for this observation.

### Effect of mupirocin resistance on virulence regulating proteins

The first virulence related protein abundance difference between the SH1000 and its mup^R^ mutant we noted was the AgrA protein^24^. This is the cytoplasmic response regulator of the Agr toxicity regulating system which was produced at significantly higher levels in the mutant (7-fold, Supp. Table 3). The *agrA* and *agrC* genes are co-transcribed, however, there was no difference in AgrC protein abundance across the protein preparations, which suggests that the effect on AgrA abundance must be post-transcriptional. The AgrA protein needs to be phosphorylated to become transcriptionally active, as such, were it active we would expect to see increased transcription of the RNAIII effector molecule of the Agr system. To test this we performed qRT-PCR on mRNA extracted from both the SH1000 and mup^R^ mutant, where we found no difference in RNAIII expression (n=6, two-tailed t-test, p=0.49), which suggests that although AgrA may be more abundant in the mutant it does not seem to be transcriptionally active.

Other known virulence regulators were also identified as being differentially produced in the mupirocin resistant strain. Both the SrrB and SarR proteins^24^ were expressed at higher levels in the mutant, whereas the Rot protein^24^ was expressed at lower levels compared to the wild type strains. Three other known regulatory proteins were also differentially expressed (i.e. a GntR family transcriptional regulator, the lytic regulatory protein SAOUHSC_02390, and the Pur operon regulator, PurR), which when considered alongside the effect on the three virulence regulators suggests that mupirocin resistance results in a significant re-wiring of *S. aureus* regulatory processes.

### Effect of mupirocin resistance on toxin production

Of the toxins encoded on the SH1000 genome, there was significantly more of both delta toxin and PSMα1 in the lysate of mupirocin resistant mutant (32- and 21-fold respectively, Supp. Table 3). These are two of the most abundantly produced members of the Phenol Soluble Modulin (PSM) family of cytolytic toxins^25^. As our original GWAS analysis was focused on toxicity, we hypothesised that these differences may explain the association we observed, and it suggests that the mutant should be more toxic than the wild type strain. To test this we quantified the cytolytic activity of the wild type and mutant by incubating bacterial supernatant with cultured THP-1 cells, which are sensitive to the majority of cytolytic toxins produced by *S. aureus*, including the PSMs^22^. While we found that mupirocin resistance did influenced toxicity, the effect was the opposite to what our proteomic analysis suggested it should be, in that the mupirocin resistant MY40 strain killed only 71% of the cells compared to the wild type mupirocin sensitive SH1000 strain which killed 87% of the cells (n=6, two tailed t-test; p=0.023). As our toxicity assays uses bacterial supernatant, whereas our proteomic analysis was on whole cell lysates, it is possible that the difference in abundance of the toxins is not equivalent between the intra- and extra-cellular environments. To examine this we quantified the abundance of PSMs in the bacterial supernatants where we found the PSMs to be 3.4-fold less abundant in the supernatant of the mup^R^ mutant (n=6, two tailed t-test; p=0.001, a representative image of is presented in fig. 2a), which explains our cytolytic results.

**Fig. 2:**
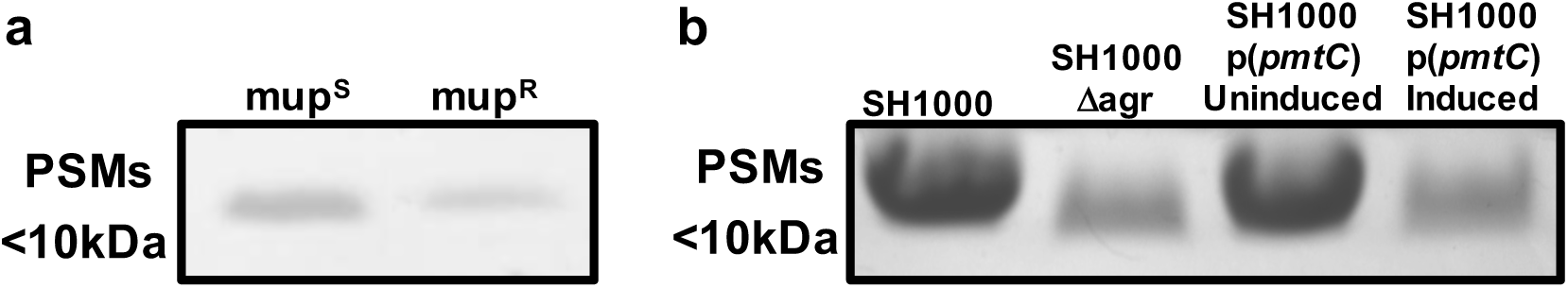
PSM abundance in the *S. aureus* supernatant. The bacterial strains were grown in TSB for 18hrs. The PSMs were harvested from the bacterial supernatant by butanol extraction and run on an SDS-PAGE gel. **a:** There is significantly less PSM in the supernatant of the mupirocin resistant *S. aureus* strain compared to the wild type mupirocin sensitive strain, **b:** Over-expression of the *pmtC* gene, which encodes one of the ATP binding proteins of the PSM secretory system, Pmt, in the wild type SH1000 strain causes a reduction in the abundance of the PSM in the *S. aureus* supernatant. An Agr mutant has been provide as a control. A full length SDS-PAGE gel has been provided in Supplementary material (Supp. Fig. 1), to illustrate why we only provide a letter-box’ snap-shot of the gels here.

### Mupirocin resistance affects the activity of the PSM secretory apparatus, Pmt

As the PSMs are more abundant in the intracellular environment of the mup^R^ mutant, but less abundant in the extracellular environment, we hypothesised that PSM secretion may be affected in the mutant. This is facilitated by the activity of the PSM export system, Pmt^26^, and although the *pmt* genes have not been annotated on the NCTC8325 genome, they are present (locus tags SAOUHSC_02155, SAOUHSC_02154, SAOUHSC_02153, SAOUHSC_02152, SAOUHSC_02151). The locus encodes a regulatory protein, two membrane bound ABC transporter proteins and two ATP binding proteins that fuel the export system, where the four structural genes are transcribed from a single promoter. Each ATP binding protein interacts with its paired transporter protein and the maintained selection of the 1:1:1:1 stoichiometry suggests it is critical to the exporters activity. In our mup^R^ mutant we found that there was more than twice as much of one of the two ATP binding proteins (SAOUHSC_02152, gene name *pmtC*) compared with the wild type strain. We hypothesised that interference with the stoichiometry of the proteins in this export system may explain why we have more PSM inside the cell but less outside. To test this we cloned and expressed the *pmtC* gene, from an inducible promoter in the SH1000 wild type background, and found that there was 2.8-fold less PSMs in the extracellular environment when this ATPase was overexpressed (n=6, two tailed t-test; p=0.003, a representative image of is presented in fig. 2b). This suggests that the effect mupirocin resistance has on *S. aureus* toxicity is mediated by interfering with the activity of the PSM secretory apparatus. It is interesting to consider that the complete inactivation of the Pmt system has been shown to be lethal to the bacteria, presumably as a result of the damage the PSMs can cause to internal membrane structures^26^. While we demonstrate a partial blocking of the Pmt system by mupirocin resistance, and some accumulation of PSMs internally, it must be a subtoxic level, as we see no *in vitro* growth defects associated with this.

### Reducing the production of toxins alleviates the fitness cost of mup^R^

With one consequence of the significant changes mup^R^ causes to the proteome of *S. aureus* being a reduction in toxin production, and with toxin production and secretion being an energetically costly activity, we hypothesized that the down regulatory effect of mupirocin resistance on toxin production may mask or alleviate the resistance related fitness costs that are incurred. To test this we quantified the relative fitness^27^ of the mup^S^ and mup^R^ strains by competition in two genetic backgrounds; one in the SH1000 wild-type background (i.e. SH1000 and MY40) where the bacteria can express toxins. For the other, we inactivated the Agr quorum sensing system in both SH1000 and MY40 by transducing in the inactivated Agr system (an erythromycin resistance has been inserted into it) from the strain ROJ48^28^, resulting in strains MY18 and MY41. As reported previously, in the wild type background with a functional Agr system, there was no difference in fitness between the mup^S^ SH1000 and the mup^R^ MY40 strains (fig. 3A; n=10; two-tailed t-test p=0.6). However, when we quantified the relative fitness in the Agr defective background where neither strain could produce toxins, such that any alleviation of fitness costs that result from the relative reduction in toxicity of the mup^R^ mutant was nullified, we found the fitness of the mup^R^ MY41 strain to be significantly lower than the mup^S^ MY18 strain (fig. 3A; n=10; two-tailed t-test p=0.04). This demonstrates that this mupirocin resistance conferring mutation does incur a fitness cost, but this cost can be masked by reducing the costly production of toxins.

**Fig. 3:**
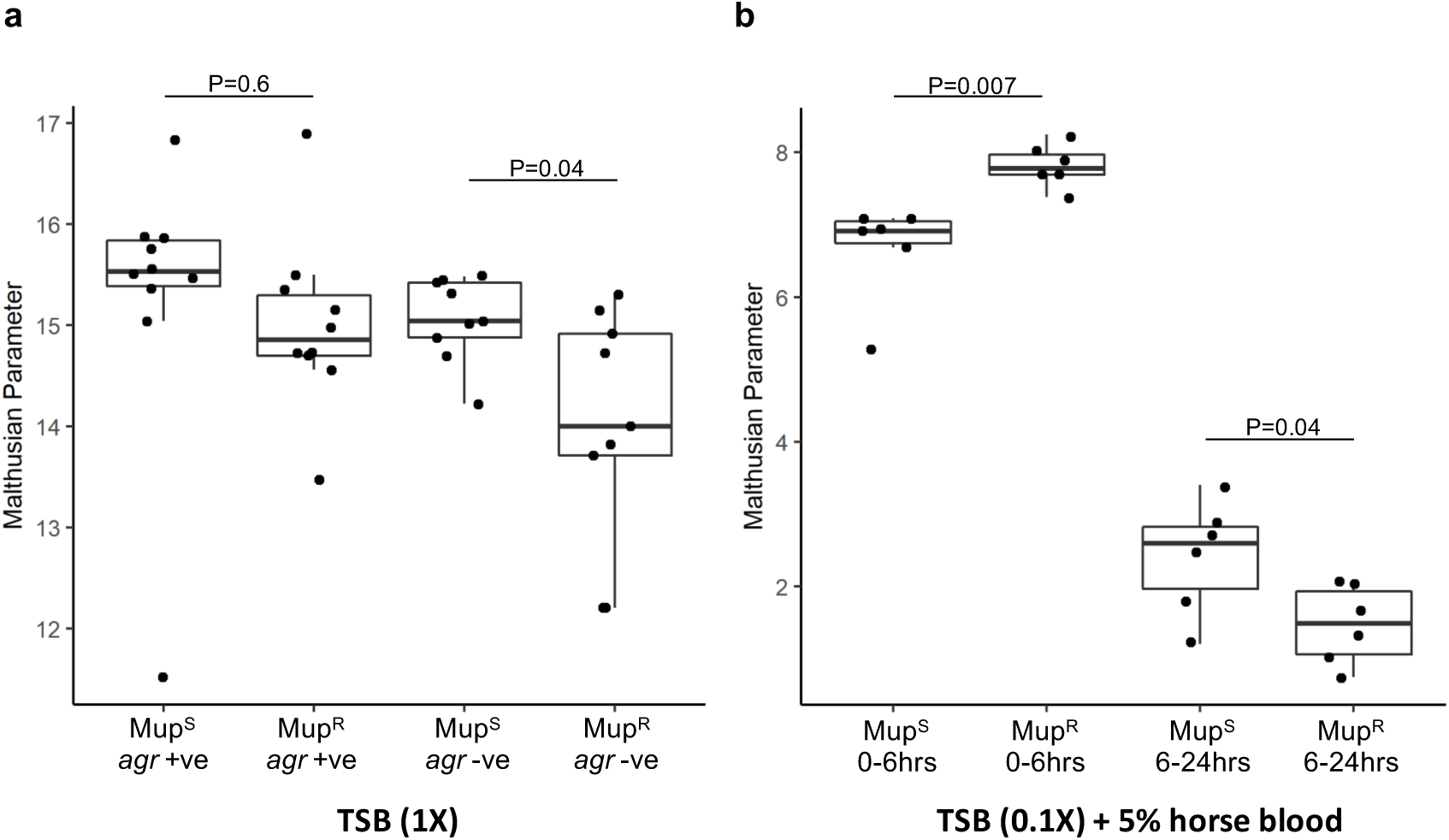
Mupirocin resistance affect the relative fitness of *S. aureus*, **a:** the effect of mupirocin resistance on the relative fitness of *S. aureus* was determined in strains with and without a functioning Agr system by direct competition in TSB. There was no difference in fitness in the Agr +ve background, but in the absence of Agr, the mupirocin resistant strain was less fit that the mupirocin sensitive strain, **b:** The effect of mupirocin resistance on relative fitness was determined by individual culture in a nutrient poor environment (0.1× TSB) supplemented with 5% horse blood. At the early stages of growth (0-6hrs) the mupirocin sensitive strain was more fit, whereas between 6 and 24hr when the nutrients in the TSB were depleted and cell lysis became necessary, the mupirocin sensitive strains was relatively more fit.

As the ability to produce toxins is selected for in environments such as the nose^29^, we hypothesised that in such an environment, the costs associated with this resistance mechanism should become apparent, without having to genetically manipulate the Agr system. To test this we developed a growth media with very low levels of nutrients (0.1 Х TSB) but added 5% horse blood such that strains that can produce toxins can release and utilise further nutrients from these blood cells for growth. To avoid the less toxic cells benefiting from nutrients the more toxic cells release, we quantified the growth rate of SH1000 and its mup^R^ mutant separately, quantifying their Malthusians parameters at 3, 6 and 24hours in this medium. After 3hr of incubation in the medium there was no detectable growth of either strain, however, by 6hrs both bacterial strains had grown, although the mup^R^ strain was significantly more fit than the SH1000 strain (fig. 3B; n=6; two-tailed t-test p=0.007). This suggests the bacteria enter a long lag phase in this medium, but once adapted (after 3hrs) there was sufficient nutrients available to sustain some growth, with the SH1000 strain at an apparent disadvantage, presumably by expending energy on producing more toxins than its mup^R^ mutant. After the 6hr time-point there was a distinct shift in the relative fitness of the strains, with the SH1000 strain becoming relatively more fit than the mup^R^ strain (fig. 3B; n=6; two-tailed t-test p=0.04). The increased relative growth rate of the wild type SH1000 strain is presumably as a result of its increased capability to lyse cells and release nutrients necessary for its growth. While the relative fitness of any organism is dependent upon its environment, here we demonstrate how readily this can fluctuate within an environment. That the fitness consequences of mupirocin resistance became apparent when cell lysis became necessary may provide an explanation for why, despite the ease at which this mutation can occur, it is not more prevalent and is readily lost within healthy communities^20^.

### The mup^R^ mutation has differing effects on toxicity in different *S. aureus* backgrounds

Our proteomic analysis did not provide any evidence in support of a direct interaction occurring between the mup^R^ *ileS* gene and the loci our GWAS analysis has identified as interacting with this locus to affect toxicity (fig. 1 and Supp. Table 1). As GWAS in bacteria can be confounded by population structures and linkage disequilibrium, it is therefore possible that these polymorphic loci are instead reflective of the genetic backgrounds of the bacteria that are sensitive to the effect of the mup^R^ mutation to varying degrees. If true, then the introduction of the mup^R^ mutation into different *S. aureus* strains should have differing effects on toxicity. To test this we isolated mup^R^ colonies of two temporally and geographically diverse isolates from the same clone as SH1000 (ST8 as determined by MLST), in the laboratory strain RN6390B and in the USA300 clinical isolate USFL34. The presence of the V588F mutation was confirmed by sequencing and the effect of the mutation on toxicity quantified. Despite reducing the toxicity of the SH1000 background strain, the mup^R^ mutation had no effect on toxicity in either RN6390B or USFL34 (n=6; two-tailed t-test p=0.59 and 0.12 respectively), supporting our hypothesis that strains respond differently to this antibiotic resistant mechanism, and providing an explanation for our epistasis findings.

## CONCLUSION

Understanding the evolution of antibiotic resistance, both in terms of how it emerges and how it becomes stably maintained, is critical if identifying means of curtailing its further emergence are to be developed. With some antibiotic resistance mechanisms being reported as incurring no fitness costs^4-6^, and others where physiological and regulatory compensation of costs have been demonstrated^30-32^, it is perhaps surprising that they have not become more prevalent. Here, by adopting a population-based functional genomics approach, we uncover hidden fitness costs associated with an apparently cost-free, antibiotic resistance mechanism. We demonstrate that the mupirocin-resistance conferring mutation of the gene encoding the IleRS enzyme has pleiotropic effects on bacteria, which in hindsight is perhaps unsurprising given the highly conserved and essential nature of this gene. In this instance, while reducing the energetic expense associated with toxin production provides a relief of the resistance-associated fitness costs, this appears to be unsustainable when the ability to produce toxins is required. Given that the nose of healthy carriers has been shown to be such a toxicity-dependent environment for *S. aureus*, this toxicity related fitness off-setting may explain why this resistance mechanism is not more prevalent, and is readily displaced, once this antibiotic has been removed from their environment. An effect we would have been unable to explain had we not adopted a population-based approach to studying this major bacterial pathogen.

## MATERIALS AND METHODS

### Strains and growth conditions

All strains used in this study are listed in Supp. Table 5. *S. aureus* strains were grown at 37°C, in either tryptic soy agar or broth (TSA/TSB) with the appropriate antibiotic where necessary. The *E. coli*TOP10 strain containing the pAgrC(his)A plasmid was grown in Luria-Bertani (LB) media with 100μg/ml ampicillin. THP-1 cells were grown in RPMI 1640 supplemented with foetal bovine serum (10%), L-glutamine (2mM), penicillin (100 units/ml) and streptomycin (100μg/ml) and incubated at 37°C with 5% CO_2_.

### Selection of the mupirocin resistant strain

As an essential gene *ileS* cannot be inactivated, and as a consequence, a homologous recombination based mutational approach were unsuccessful. Therefore, to generate isogenic mupirocin resistant and sensitive strains we utilised a selection based method where an overnight culture of a mupirocin sensitive strain (e.g SH1000 was plated onto agar plates with 4μg/ml mupirocin. This was incubated at 37°C for 48 hr and colonies that grew were further isolated by streaking onto fresh mupirocin plates.

### Genome sequencing of the mup^R^ strain

*S. aureus* strain MY40 was sequenced in this study; DNA was extracted using the QIAamp DNA Mini Kit (QIAGEN, Crawley, UK), using manufacturer’s instructions with 1.5 μg/μL lysostaphin (Ambi Products LLC, NY, USA) to facilitate cell lysis. DNA was quantified using a Nanodrop spectrophotometer, as well as the Quant-iT DNA Assay Kit (Life Technologies, Paisley, UK) before sequencing. High-throughput genome sequencing was performed using a MiSeq machine (Illumina, San Diego, CA, USA) and the short read, paired-end data was assembled using the *de novo* assembly algorithm SPAdes (Bankevich et al (SPADES). Sequence data are archived in the NCBI repositories: GenBank Accession: SUB2754769, Short Read Archive (SRA): SRR5651527, associated with BioProject: PRJNA384009. Assembled genomes are also available on FigShare (doi.org/10.6084/m9.figshare.5089939.v1).

### Toxicity assays

THP-1 cells^33^ were grown as described above and harvested by centrifugation and washed in PBS and diluted to a final density (determined by haemocytometer) of 2×10^6^ cells per ml of PBS. Bacterial supernatant was harvested after 18hrs of growth in TSB at 37°C. 20μl of the bacterial supernatant was mixed with 20μl of THP-1 cells, and incubated for 12 mins at 37°C. 260μl of Guava ViaCount (Milipore) was added to the sample, and incubated at room temperature for 5 mins before analysing the viability on the Guava flow cytometer (Milipore).

### Fitness assays

The strains were cultured individually overnight and diluted to 10^4^ cfu/ml. For the direct competitions 25μl of each diluted culture was added into 5ml fresh TSB, and grown at 37°C with shaking for 24h. The mixed culture was diluted and plated onto agar plates with and without 4μg/ml mupirocin and incubated at 37°C. The resulting colonies were counted, and the number of colonies from the mupirocin plate was subtracted from the count from no antibiotic plate. The Malthusian parameter was calculated using the following formula:

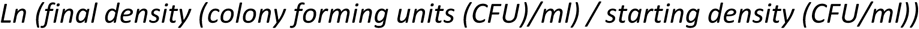

The Malthusian parameters of the mup^S^ and mup^R^ strains were compared using a two tailed t-test. For the individual fitness assays the strains were cultured individually overnight and diluted and added to 0.1× TSB made with phosphate buffered saline rather than water to maintain the integrity of the horse blood cells to which and 5% horse blood (Oxoid) was added. Bacterial growth was determined by plating the cultures on TSA plates after 3, 6 and 24hrs of incubation at 37°C in air.

### Deletion of the *agr* locus from SH1000 and MY40

Phage transduction was used to construct the mup^R^ and mup^S^ Agr mutants. In the *S. aureus* strain ROJ48, the entire Agr locus has been replaced with an erythromycin resistance cassette^28^. This was moved from ROJ48 into SH1000 by phage transduction as follows: ROJ48 φ11 lysates were prepared from 200μl of overnight ROJ48 culture in LK (1% Tryptone, 0.5% yeast extract, 1.6% KCl) which was added to 3ml of fresh LK and 3ml of phage buffer (10mM MgSO_4_, 4mM CaCl_2_, 50mM Tris-HCl pH 7.8, 100mM NaCl and 0.1% gelatine powder in molecular/MiliQ water), and to this 500μl φ11-RN6390B lysate was added. This was incubated at 30°C shaking until the media became clear, which indicated bacterial lysis. The lysates were then filter sterilised, and a second round of lysis was carried out on ROJ48 with this first round lysate. After these two lysis steps, transduction into SH1000 and MY40 strains was performed by adding 200μl of overnight culture to 1.8ml LK with 10μl 1M CaCl_2_, and 500μl the φ11-ROJ48 lysate. This was incubated at 37°C with shaking for 45 min, then 1ml ice cold 20mM trisodium citrate was added and the transducing mixture was placed on ice for 5 min. The bacteria were harvested by centrifugation and resuspended with 1ml ice cold 20mM trisodium citrate. This was incubated on ice for 2.5h, and plated onto TSA plates with 20mM trisodium citrate, erythromycin and lincomycin (25μg/ml) which was incubated overnight at 37°C.

### qRT-PCR

Overnight cultures of were diluted 1:500 into 3ml fresh TSB-chloramphenicol. After 18hr of growth 2 ml of this culture was mixed with 4ml RNA Protect Bacteria (Qiagen), and the RNeasy Mini Kit (Qiagen) was used to extract RNA following the manufacturer’s protocol. Lysostaphin (200μg/ml) was added to Tris-EDTA buffer (Ambion), and this was added to the sample after the RNA Protect step before continuing with the protocol. When the RNA was extracted, Turbo DNA-free kit (Thermo) was used to remove genomic DNA from the RNA samples; 3μl Turbo DNase was added to the sample and incubated for 1.5h at 37°C, the a further 4μl Turbo DNase was added and incubated for 1.5h. 35μl DNase inactivation reagent was added to the samples to inactivate the DNase according to the protocol. The concentration of RNA in the samples were measured using Qubit RNA Broad Range kit (Thermo) and normalised before using QuantiTect Reverse Transcription Kit (Qiagen) to convert the RNA samples into cDNA according to the manufacturer’s protocol. After adding the reverse transcriptase, the samples were incubated at 42°C for 20 min before raising the temperature to 95°C for 3 mins to inactivate the reverse transcriptase. Primers for *gyrB*, a housekeeping gene, was used alongside those for RNAIII to standardise transcript levels (gyrB forward: CCAGGTAAATTAGCCGATTGC, gyrB reverse: AAATCGCCTGCGTTCTAGAG. RNAIII forward; AGCATGTAAGCTATCGTAAACAAC, RNAIII reverse; TTCAATCTATTTTTGGGGATG). ssoAdvanced SYBR Green Supermix (Bio-Rad) was used, using a standard curve of known genomic DNA concentrations for each primer set. 5μl of samples, standards and water were pipetted into the wells of a 96-well PCR plate. The supermix was added to water and primers according to the manufacturer’s protocol, and 15μl of this mix was pipetted over the DNA samples. This was then placed into a qPCR machine, and run using the manufacturer’s recommendation. The quantity of RNA III cDNA was divided by the quantity of *gyrB* cDNA to get a ratio of RNAIII transcription levels.

### PSM quantification in supernatants

An overnight culture of SH1000 and MY40 was diluted 1:1000 into 50ml fresh TSB, and grown for 18h. The cultures were centrifuged at 18,000 rpm for 10 min, and 35ml of the supernatant was mixed with 10ml butanol. The samples were incubated at 37°C shaking for 3h, and were then centrifuged at 3,000 rpm for 3 min and 1ml of the upper organic layer was taken off. The samples were then freeze-dried overnight and then re-suspended in 160μl 8M urea, separated on an SDS-PAGE gel and PSMs quantified by densitometry analysis using the ImageJ software.

### TMT Labelling and High pH reversed-phase chromatography

Aliquots of 100μg of up to ten samples per experiment were digested with trypsin (2.5μg trypsin per 100μg protein; 37°C, overnight), labelled with Tandem Mass Tag (TMT) ten plex reagents according to the manufacturer’s protocol (Thermo Fisher Scientific,) and the labelled samples pooled. An aliquot of the pooled sample was evaporated to dryness and resuspended in buffer A (20 mM ammonium hydroxide, pH 10) prior to fractionation by high pH reversed-phase chromatography using an Ultimate 3000 liquid chromatography system (Thermo Fisher Scientific). In brief, the sample was loaded onto an XBridge BEH C18 Column (130Å, 3.5 μm, 2.1 mm × 150 mm, Waters, UK) in buffer A and peptides eluted with an increasing gradient of buffer B (20 mM Ammonium Hydroxide in acetonitrile, pH 10) from 0-95% over 60 minutes. The resulting fractions were evaporated to dryness and resuspended in 1% formic acid prior to analysis by nano-LC MSMS using an Orbitrap Fusion Tribrid mass spectrometer (Thermo Scientific).

### Nano-LC Mass Spectrometry

High pH RP fractions were further fractionated using an Ultimate 3000 nanoHPLC system in line with an Orbitrap Fusion Tribrid mass spectrometer (Thermo Scientific). In brief, peptides in 1% (vol/vol) formic acid were injected onto an Acclaim PepMap C18 nano-trap column (Thermo Scientific). After washing with 0.5% (vol/vol) acetonitrile 0.1% (vol/vol) formic acid peptides were resolved on a 250 mm × 75 μm Acclaim PepMap C18 reverse phase analytical column (Thermo Scientific) over a 150 min organic gradient, using 7 gradient segments (1-6% solvent B over 1min., 6-15% B over 58min., 15-32%B over 58min., 32-40%B over 5min., 40-90%B over 1min., held at 90%B for 6min and then reduced to 1%B over 1min.) with a flow rate of 300 nl min^-1^. Solvent A was 0.1% formic acid and Solvent B was aqueous 80% acetonitrile in 0.1% formic acid. Peptides were ionized by nano-electrospray ionization at 2.0kV using a stainless steel emitter with an internal diameter of 30 μm (Thermo Scientific) and a capillary temperature of 275°C.

All spectra were acquired using an Orbitrap Fusion Tribrid mass spectrometer controlled by Xcalibur 2.0 software (Thermo Scientific) and operated in data-dependent acquisition mode using an SPS-MS3 workflow. FTMS1 spectra were collected at a resolution of 120 000, with an automatic gain control (AGC) target of 200 000 and a max injection time of 50ms. Precursors were filtered with an intensity threshold of 5000, according to charge state (to include charge states 2-7) and with monoisotopic precursor selection. Previously interrogated precursors were excluded using a dynamic window (60s +/-10ppm). The MS2 precursors were isolated with a quadrupole mass filter set to a width of 1.2m/z. ITMS2 spectra were collected with an AGC target of 10 000, max injection time of 70ms and CID collision energy of 35%. For FTMS3 analysis, the Orbitrap was operated at 50 000 resolution with an AGC target of 50 000 and a max injection time of 105ms. Precursors were fragmented by high energy collision dissociation (HCD) at a normalised collision energy of 60% to ensure maximal TMT reporter ion yield. Synchronous Precursor Selection (SPS) was enabled to include up to 5 MS2 fragment ions in the FTMS3 scan.

### Proteomic data analysis

The raw data files were processed and quantified using Proteome Discoverer software v2.1 (Thermo Scientific) and searched against the UniProt Staphylococcus aureus strain NCTC 8325 database using the SEQUEST algorithm. Peptide precursor mass tolerance was set at 10ppm, and MS/MS tolerance was set at 0.6Da. Search criteria included oxidation of methionine (+15.9949) as a variable modification and carbamidomethylation of cysteine (+57.0214) and the addition of the TMT mass tag (+229.163) to peptide N-termini and lysine as fixed modifications. Searches were performed with full tryptic digestion and a maximum of 2 missed cleavages were allowed. The reverse database search option was enabled and all peptide data was filtered to satisfy false discovery rate (FDR) of 5%.

## REFERENCES

1. Moura de Sousa, J., Balbontín, R., Durão, P. & Gordo, I. Multidrug-resistant bacteria compensate for the epistasis between resistances. PLOS Biol. 15, e2001741 (2017).

2. Tenover, F. C. Mechanisms of Antimicrobial Resistance in Bacteria. Am. J. Med. 119, S3–S10 (2006).

3. Levin, B. R., Perrot, V. & Walker, N. Compensatory mutations, antibiotic resistance and the population genetics of adaptive evolution in bacteria. Genetics 154, 985–997 (2000).

4. Hurdle, J. G., O’Neil, A. J. & Chopra, I. The isoleucyl-tRNA synthetase mutation V588F conferring mupirocin resistance in glycopeptide-intermediate Staphylococcus aureus is not associated with a significant fitness burden. J. Antimicrob. Chemother. 53, 102–104 (2003).

5. Stanczak-Mrozek, K. I. et al. Within-host diversity of MRSA antimicrobial resistances. J. Antimicrob. Chemother. 70, 2191–2198 (2015).

6. Knight, G. M., Budd, E. L. & Lindsay, J. A. Large mobile genetic elements carrying resistance genes that do not confer a fitness burden in healthcare-associated meticillin-resistant Staphylococcus aureus. Microbiol. (United Kingdom) 159, 1661–1672 (2013).

7. Rasigade, J.-P. & Vandenesch, F. Staphylococcus aureus: a pathogen with still unresolved issues. Infect. Genet. Evol. 21, 510–4 (2014).

8. Lowy, F. Antimicrobial resistance: the example of Staphylococcus aureus. J. Clin. Invest. 111, 1265–1273 (2003).

9. Wertheim, H. F. et al. The role of nasal carriage in Staphylococcus aureus infections. Lancet Infect. Dis. 5, 751–762 (2005).

10. Gordon, R. J. & Lowy, F. D. Pathogenesis of Methicillin - Resistant Staphylococcus aureus Infection. Clin. Infect. Dis. 46, S350–S359 (2008).

11. Abad, C. L., Pulia, M. S. & Safdar, N. Does the Nose Know? An Update on MRSA Decolonization Strategies. Curr. Infect. Dis. Rep. 15, 455–464 (2013).

12. Thomas, C. M., Hothersall, J., Willis, C. L. & Simpson, T. J. Resistance to and synthesis of the antibiotic mupirocin. Nat. Rev. Microbiol. 8, 281–289 (2010).

13. Simor, A. E. Staphylococcal decolonisation: An effective strategy for prevention of infection? Lancet Infect. Dis. 11, 952–962 (2011).

14. Bertino Jr, J. S. Intranasal mupirocin for outbreaks of methicillin-resistant Staphylococcus aureus. Am J Heal. Pharm 54, 2185–2191 (1997).

15. Antonio, M., McFerran, N. & Pallen, M. J. Mutations affecting the Rossman fold of isoleucyl-tRNA synthetase are correlated with low-level mupirocin resistance in Staphylococcus aureus. Antimicrob. Agents Chemother. 46, 438–442 (2002).

16. Patel, J. B., Gorwitz, R. J. & Jernigan, J. A. Mupirocin Resistance. Clin. Infect. Dis. 49, 935–941 (2009).

17. Udo, E. E. & Sarkhoo, E. Genetic analysis of high-level mupirocin resistance in the ST80 clone of community-associated meticillin-resistant Staphylococcus aureus. J. Med. Microbiol. 59, 193–199 (2010).

18. Seah, C. et al. MupB, a New High-Level Mupirocin Resistance Mechanism in Staphylococcus aureus. Antimicrob. Agents Chemother. 56, 1916–1920 (2012).

19. Hetem, D. J. & Bonten, M. J. M. Clinical relevance of mupirocin resistance in Staphylococcus aureus. J. Hosp. Infect. 85, 249–256 (2013).

20. Williamson D.A., Carter G.P. & Howden B.P. Current and Emerging Topical Antibacterials and Antiseptics: Agents, Action, and Resistance Patterns. Clin. Microbiol. Rev. 30, 827-860 (2017).

21. Laabei, M. et al. Predicting the virulence of MRSA from its genome sequence. Genome Res. 24, 839–849 (2014).

22. Laabei et al. Evolutionary Trade-Offs Underlie the Multi-faceted Virulence of Staphylococcus aureus. PLoS Biol. 13(9):e1002229 (2015).

23. Dayon L & JC Sanchez. Relative protein quantification by MS/MS using the tandem mass tag technology. Methods Mol Biol. 893:115-27 (2012).

24. Bronner, S., Monteil, H. & Prévost, G. Regulation of virulence determinants in Staphylococcus aureus: complexity and applications. FEMS Microbiol. Rev. 28, 183–200 (2004).

25. Qi, R. et al. Increased In Vitro Phenol-Soluble Modulin Production is Associated with Soft Tissue Infection Source in Clinical Isolates of Methicillin-Susceptible Staphylococcus aureus. J. Infect. 72, 302–308 (2017).

26. Chatterjee, S. S. et al. Essential Staphylococcus aureus toxin export system. Nat. Med. 19, 364–367 (2013).

27. Lenski, R. E., Rose, M. R., Simpson, S. C. & Tadler, S. C. Long-Term Experimental Evolution in Escherichia coli. I. Adaptation and Divergence During 2,000 Generations. Am. Nat. 138, 1315–1341 (1991).

28. Jensen, R. O., Winzer, K., Clarke, S. R., Chan, W. C. & Williams, P. Differential Recognition of Staphylococcus aureus Quorum-Sensing Signals Depends on Both Extracellular Loops 1 and 2 of the Transmembrane Sensor AgrC. J. Mol. Biol. 381, 300–309 (2008).

29. Recker M, et al. Clonal differences in Staphylococcus aureus bacteraemia-associated mortality. Nat Microbiol. 2, 1381-1388 (2017).

30. Pacheco, JO, Alvarez-Ortega, C, Rico, MA & Martínez, JL. Metabolic Compensation of Fitness Costs Is a General Outcome for Antibiotic-Resistant Pseudomonas aeruginosa Mutants Overexpressing Efflux Pumps. mBio. 8, e00500-17 (2017).

31. Freihofer P, et al. Nonmutational compensation of the fitness cost of antibiotic resistance in mycobacteria by overexpression of tlyA rRNA methylase. RNA. 22, 1836-1843 (2016).

32. Collins, J et al. Offsetting virulence and antibiotic resistance costs by MRSA. ISME J. 4, 577-84 (2010).

33. Tsuchiya, S. et al. Establishment and characterization of a human acute monocytic leukemia cell line (THP - 1). Int. J. Cancer 26, 171-176 (1980).

